## Supplemental Data for "Interneuron FGF13 regulates seizure susceptibility via a sodium channel-independent mechanism"

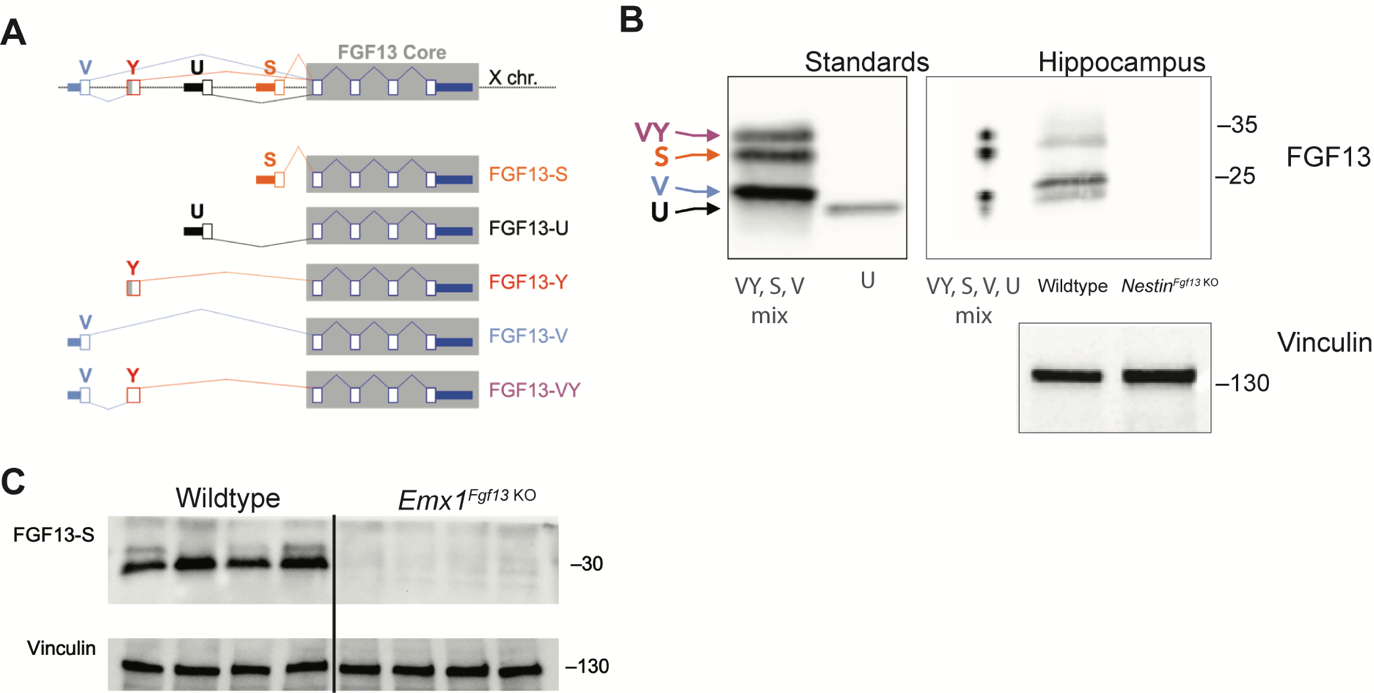

**Supplemental Figure 1. *Fgf13* splice variants are differentially expressed in hippocampal cell types.**

**A**. *Fgf13* generates five alternatively spliced variants. **B**. Knockout of *Fgf13* in *Nestin* neurons eliminates all isoforms. Standards are lysates from HEK293 cells transfected with cDNAs expressing a mix of *Fgf13-VY, -S,* and *-V* or expressing *Fgf13-U.* Western blot performed with pan-FGF13 antibody. Vinculin is used as loading control. **C**. Excitatory neuronal knockout of *Fgf13* results in loss of FGF13-S. Western blot performed with FGF13-S specific antibody. Vinculin used as a loading control.

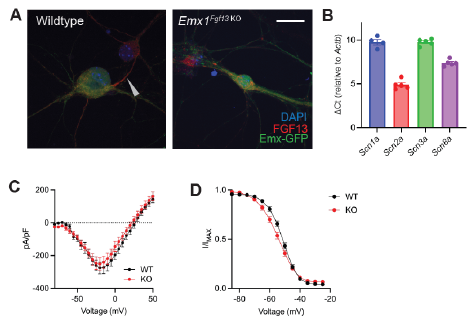

**Supplemental Figure 2. Excitatory neuronal knockout of *Fgf13* does not result in sodium current deficits.**

**A**. Primary hippocampal neuron cultures were generated from *Emx1*-Cre wildtype (left) or *Emx1^Fgf13 KO^* mice, and *Emx1*+ excitatory cells were labeled by infection with a Cre-dependent AAV8-DIO-GFP virus. Immunocytochemistry was performed with a pan-FGF13 antibody (scale bar, 20 μm). Arrow indicates FGF13 in the axon initial segment, absent in the *Emx1^Fgf13 KO^* neurons. **B**. Reverse transcriptase quantitative polymerase chain reaction of neuronal voltage-gated sodium channels shows levels of expression for specific channels relative to *Actb* (N=4-5 mice). **C**. *Emx1^Fgf13 KO^* neurons do not exhibit differences in macroscopic sodium currents (two-way ANOVA, p=ns). Peak current-voltage (I-V) curves shown; WT *N=*3 mice, n=18 cells; KO *N*=4 mice, n=18 cells. **D**. *Emx1^Fgf13 KO^* neurons do not have a significant difference in steady-state inactivation (V_1/2_ WT = -52.04 [95% CI, -53.60 to -50.47]; V_1/2_ KO = -54.47 [95% CI, -56.4 to -52.54]). WT *N=*3 n=21; KO *N*=4 n=37.

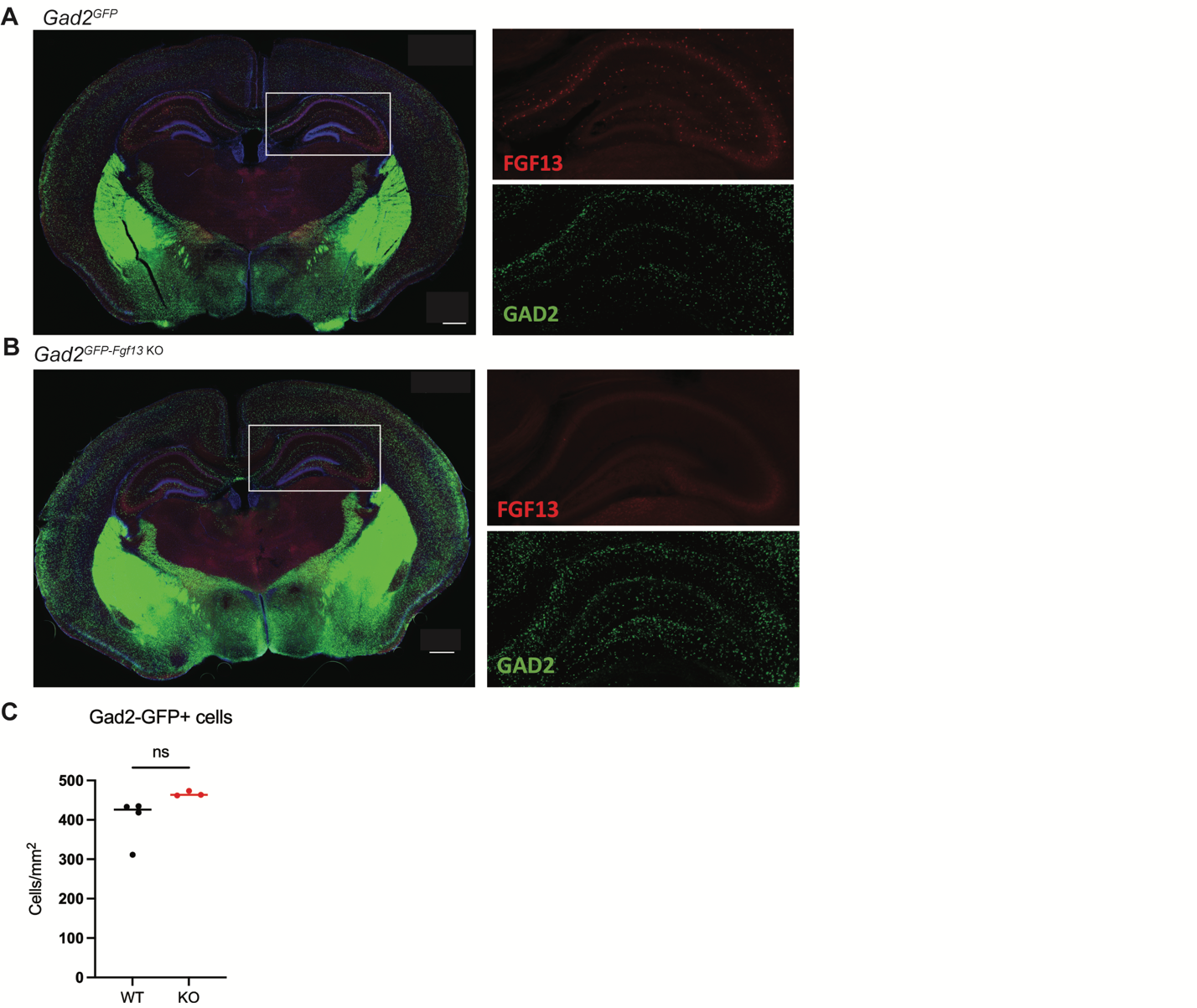

**Supplemental Figure 3. Characterization of FGF13 abundance in hippocampus of *Gad2^Fgf13 KO^* mice.**

**A**. Fluorescent immunohistochemistry with a pan-FGF13 antibody on hippocampal tissue in *Gad2^GFP^* reporter mice. **B**. Immunohistochemistry of hippocampal tissue in *Gad2^GFP-Fgf13 KO^* shows loss of FGF13 but not loss of *Gad2-*GFP^+^ interneurons (scale bar, 500 μm) and no gross morphological changes. **C**. *Gad2^Fgf13 KO^* mice do not have a significant change in the density of hippocampal GABAergic interneurons, despite decreased brain and body size relative to wildtype littermates (t-test, p=ns), WT *N*=4 mice, KO *N*=3 mice.

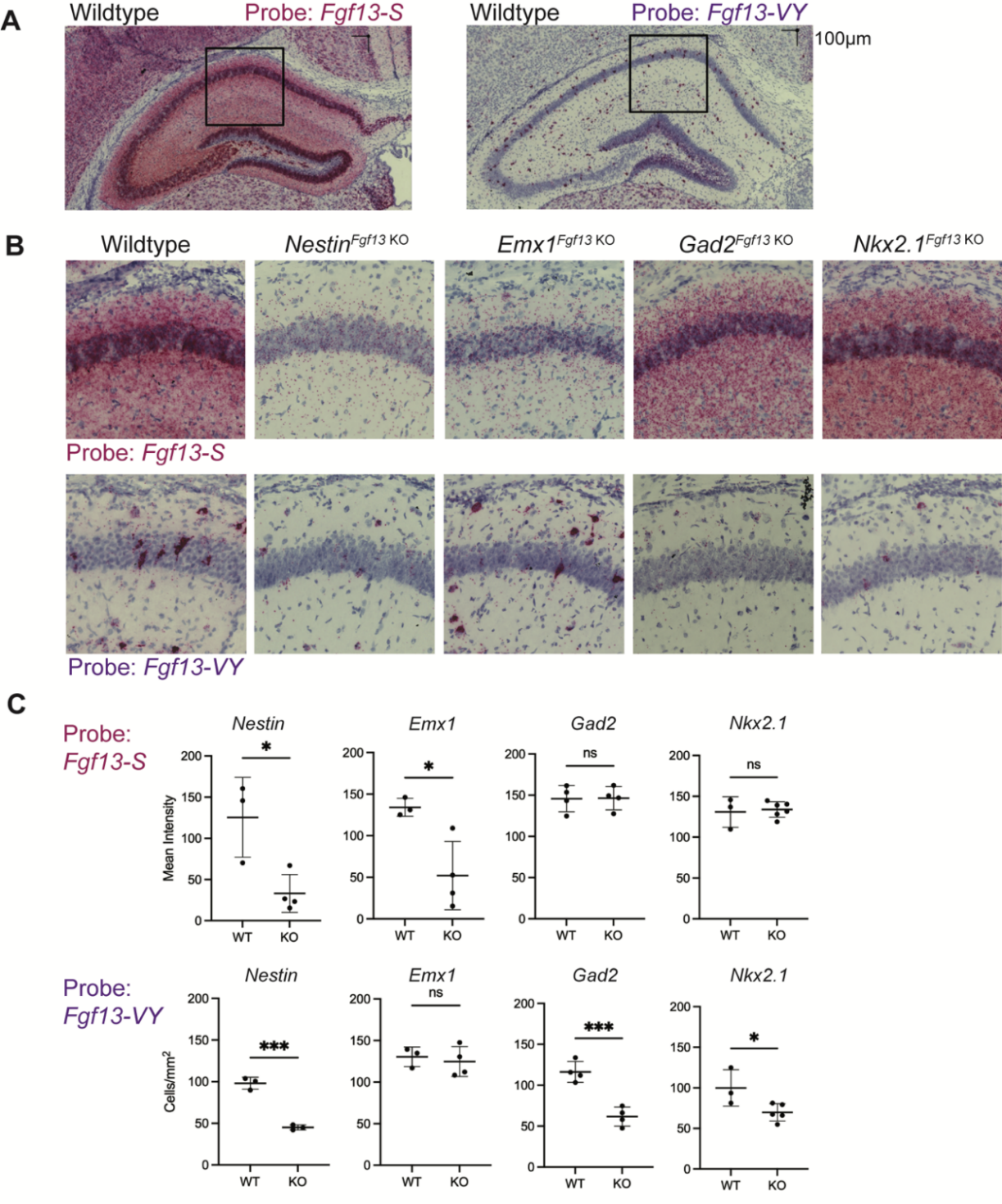
**Supplemental Figure 4. *Fgf13* splice variants are differentially expressed in hippocampal cell types.**

**A**. *Fgf13-S* and *-VY* exhibit distinct expression in the hippocampus of wildtype and cell type-specific knockouts. **B**. Example images showing decreased *Fgf13-S* expression from *Nestin^Fgf13 KO^* and *Emx1^Fgf13 KO^* hippocampus or decreased *Fgf13-VY* expression from *Nestin^Fgf13 KO^*, *Gad2^Fgf13 KO^*, and *Nkx2.1^Fgf13 KO^* hippocampus. **C**. Quantification of BASEscope *in situ* hybridization reveals that *Fgf13-S* expression is decreased in *Nestin^Fgf13 KO^* and *Emx1^Fgf13 KO^* hippocampus, while *Fgf13-VY* expressing cells are decreased in *Nestin^Fgf13 KO^*, *Gad2^Fgf13 KO^*, and *Nkx2.1^Fgf13 KO^* hippocampus. (t-test, *, p<0.05; ***, p<0.001) *N=*3-6 mice each.

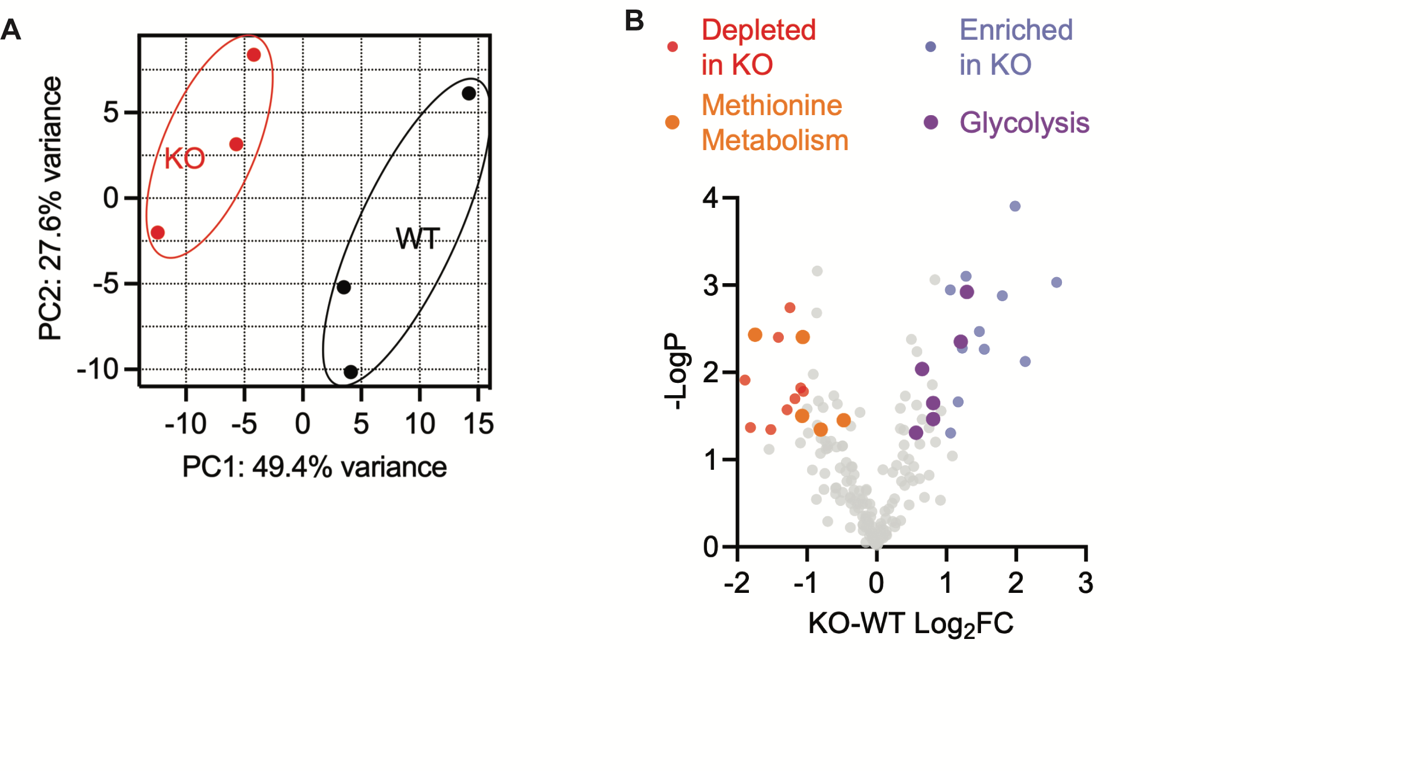

**Supplemental Figure 5. An unbiased metabolomic analysis on whole brain lysate from *Gad2^Fgf13 KO^* and wildtype mice reveals marked differences between genotypes.**

**A**. PCA plot showing 3 WT and 3 *Gad2 ^Fgf13 KO^* mice. **B**. Volcano plot of measured metabolites. Colored data show metabolites with *P*<0.05 and Log2FC > 1 or Log2FC <1 (KO v WT). *N=*3 mice each.

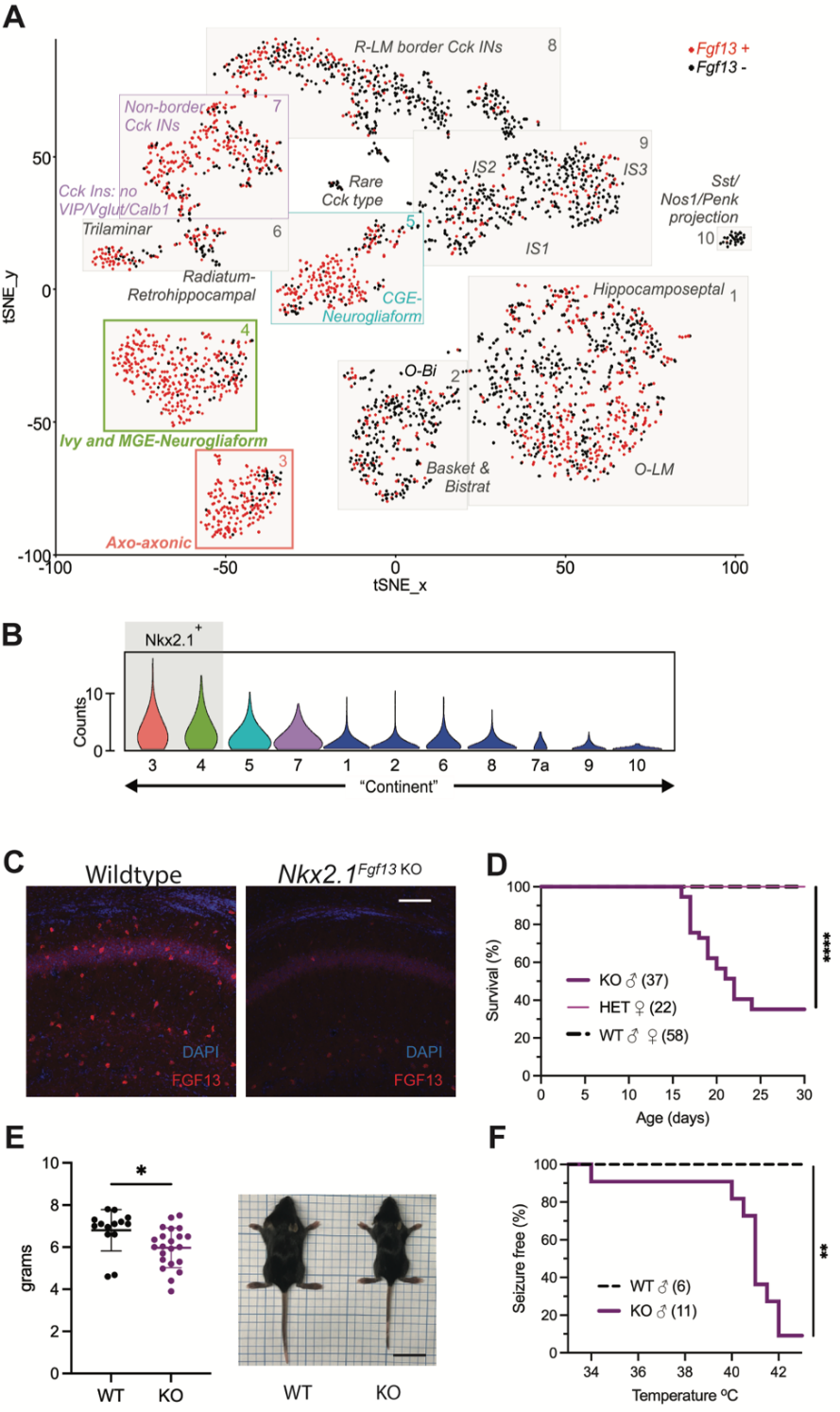

**Supplemental Figure 6. Knockout of *Fgf13* from MGE-derived interneurons recapitulates premature death and seizure susceptibility.**

**A**. Re-analysis of single-cell RNA-seq data (1) reveals that *Fgf13* is expressed in MGE-derived interneurons such as the axo-axonic and MGE-derived neurogliaform cell types, as well as CGE-derived interneurons. **B.** Violin plots show that cell type continents with the highest expression of *Fgf13^+^* cells are *Nkx2.1*^+^. **C**. Fluorescent immunohistochemistry of hippocampal tissue validates *Fgf13* knockout (scale bar, 100 μm). **D***. Nkx2.1^Fgf13 KO^* mice have survival deficits around one month of age (log-rank test, ****, p<0.0001). **E**. Body mass at postnatal day 14 (P14) shows *Nkx2.1^Fgf13 KO^* are smaller in size (t-test, *, p<0.05). Scale, 2 cm. **F**. *Nkx2.1^Fgf13 KO^* mice are susceptible to hyperthermia induced seizures (log-rank test, **, p<0.01).

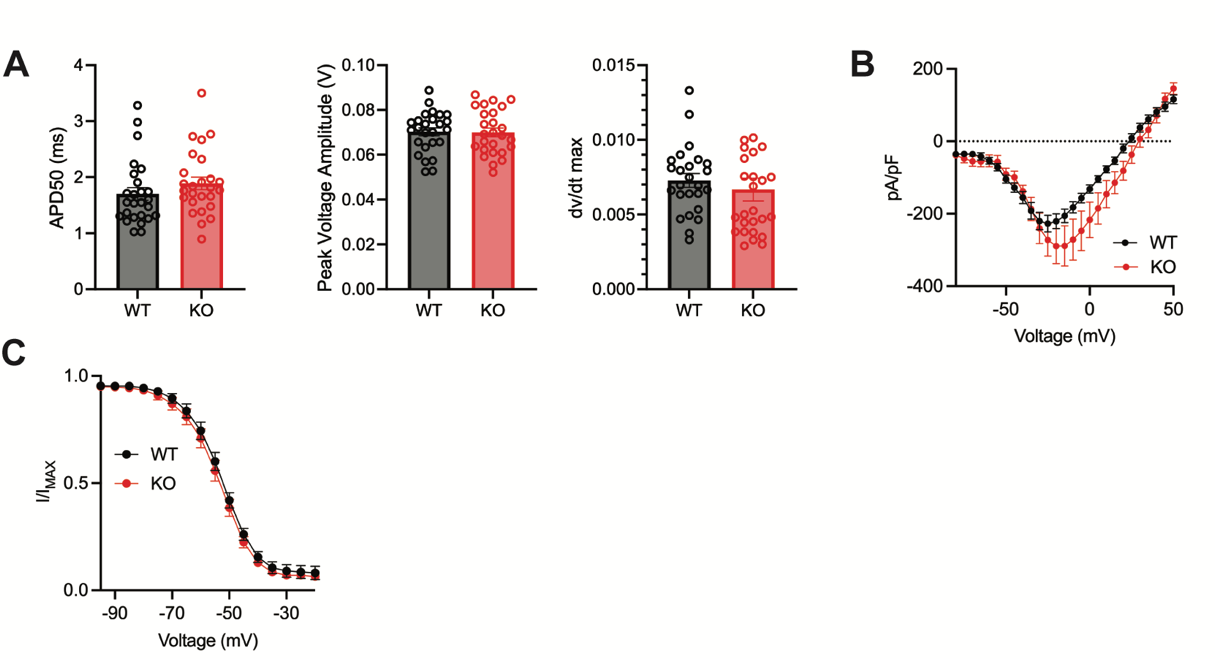

**Supplemental Figure 7. Inhibitory neuronal knockout of *Fgf13* does not affect action potential parameters nor sodium channel currents.**

**A**. Wildtype and *Gad2^Fgf13 KO^* interneurons have no significant difference in APD50, peak voltage amplitude, and dV/dt max values for the first action potential in each spike train. **B**. Wildtype and *Gad2^Fgf13 KO^* interneurons have no difference in macroscopic sodium currents (two-way ANOVA, p=ns). Peak current-voltage curves shown for WT and KO, WT *N=*3 n=25; KO *N*=3 n=18. **C**. Wildtype and *Gad2^Fgf13 KO^* interneurons have no difference in steady-state inactivation (V_1/2_ WT = -52.98 [95% CI, -55.32 to -50.63]; V_1/2_ KO = -54.19 [95% CI, -57.28 to -51.10], p = ns. *N*(WT)*=*3 n=21, N(KO)=3 n=17.

| **Supplemental Table 1: Primers used for real time quantitative polymerase chain reaction** | | | |
| --- | --- | --- | --- |
| **Forward (5’-3’)** | **Reverse (5’-3’)** | **Gene** | **PrimerBank* ID** |
| TCAGAGGGAAGCACAGTAGAC | TTCCACGCTGATTTGACAGCA | *Scn1a* | 9055328a1 |
| TTCATGGCTTCCAATCCCTCC | GGTGTCACGTCAGTCTTCTCT | *Scn2a* | 26328015a1 |
| CAGACCATGTGCCTTATTGTGT | CCGCGATCTGGAGGTTGTT | *Scn3a* | 9055330a1 |
| ATGGGGTAGGCTCTCCGAG | CCGACTCTGACTTAAACACCTTC | *Scn8a* | 6755410a1 |
| GATCAAGATCATTGCTCCTCCTG | AGGGTGTAAAACGCAGCTCA | *Actb* |  |
| *References for PrimerBank:  Spandidos A, et al. A comprehensive collection of experimentally validated primers for polymerase chain reaction quantitation of murine transcript abundance. *BMC Genomics*. 2008;9:633.  Spandidos A, et al. PrimerBank: a resource of human and mouse PCR primer pairs for gene expression detection and quantification. *Nucleic Acids Res*. 2010;38(database issue):D792–D799 | | | |

**References(2, 3)**

1.	Harris, K. D., Hochgerner, H., Skene, N. G., Magno, L., Katona, L., Bengtsson Gonzales, C. *et al.* (2018) Classes and continua of hippocampal CA1 inhibitory neurons revealed by single-cell transcriptomics PLoS Biol **16**, e2006387 10.1371/journal.pbio.2006387

2.	Spandidos, A., Wang, X., Wang, H., andSeed, B. (2010) PrimerBank: a resource of human and mouse PCR primer pairs for gene expression detection and quantification Nucleic Acids Res **38**, D792-799 10.1093/nar/gkp1005

3.	Spandidos, A., Wang, X., Wang, H., Dragnev, S., Thurber, T., andSeed, B. (2008) A comprehensive collection of experimentally validated primers for Polymerase Chain Reaction quantitation of murine transcript abundance BMC Genomics **9**, 633 10.1186/1471-2164-9-633
